## Supplementary material for "Two-way Dispatched function in Sonic hedgehog shedding and transfer to high-density lipoproteins": Contains 9 Supplemental figures and 3 Tables

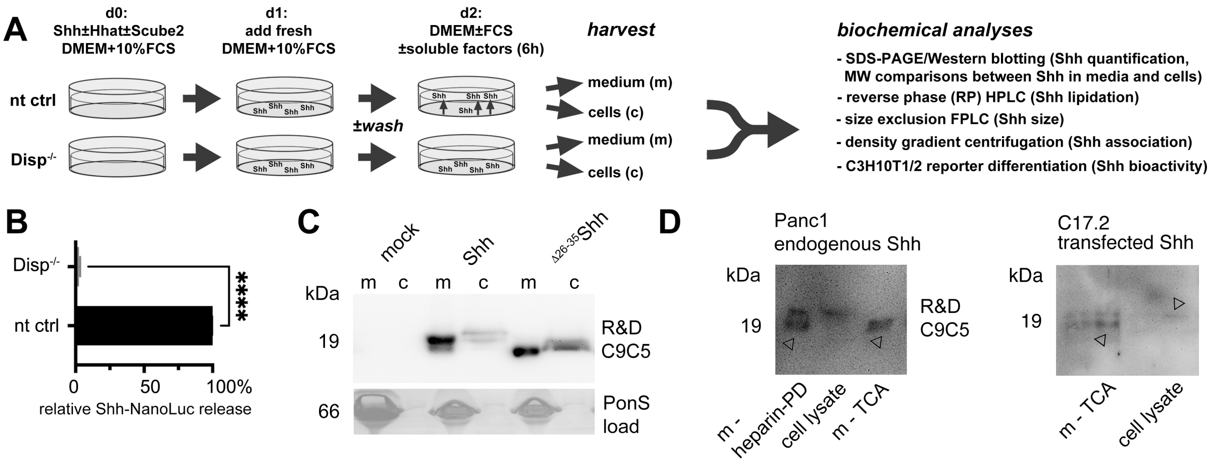


**Figure Supplement 1. Experimental design and controls. A)** nt Bosc23 ctrl cells that express Disp (top) or $Disp^{-/-}$ cells (bottom) were transfected on day 0 with Shh constructs together with Scube2 or empty control cDNA3.1. Fresh medium was added 1 day after transfection and cells were grown for 36 h (100 µg/mL) in DMEM containing 10% FCS and penicillin-streptomycin. For serum-depleted solubilization, on day 2, serum-containing media were aspirated and serum-free DMEM (with or without CMK furin inhibitor, serum, or HDL/LDL) was added for 6 h. Media were harvested and centrifuged at 300*g* for 10 min to remove debris. Supernatants containing FCS were subjected to heparin-pulldown, and serum-free or serum-depleted supernatants were incubated with 10% trichloroacetic acid for 30 min on ice, followed by centrifugation at 13,000*g* for 20 min to precipitate the proteins. Cells were lysed directly on the plate for subsequent RP-HPLC or SDS-PAGE analysis. SEC, density gradient centrifugation and cell-based Shh activity/qPCR assays were also performed. **B)** We confirmed impaired Shh solubilization from our $Disp^{-/-}$ cell line by using a Shh variant internally tagged with NanoLuciferase (Wang et al., 2021) and co-expressed with Hhat from bicistronic mRNA. Strongly reduced luminescence was detected in media from Shh-NanoLuc expressing $Disp^{-/-}$ cells. Unpaired t-test, two-tailed. *: p=0.017, n=3. Note that quantification of Shh-NanoLuc in media does not provide any information beyond increased/decreased protein release. In particular, it does not provide direct information on how Disp drives the release of dual-lipidated Hh from the plasma membrane, whether Disp acts directly or indirectly in this process, whether Shh remains fully lipidated during release, and to which carrier – if any – Hh is transferred. In contrast, the methods used in our work provide this information. **C)** Cells were transfected with full-length $^{C25S}$Shh or N-truncated $^{\Delta 26\text{-}35}$Shh, and proteins were concentrated by TCA precipitation. The C9C5 antibody used for Shh detection throughout this work proved to be highly specific. **D)** Reduced electrophoretic mobility of endogenous Shh from pancreatic ductal carcinoma (Panc1) cells (cell lysate) if compared to soluble proteins (m, arrowheads) indicates proteolytic processing of the endogenous protein during release. m-heparin-PD: Shh pulldown from serum-containing medium with heparin, m-TCA: TCA precipitation of Shh secreted into serum-depleted media. Shh was also released in processed form from transfected C17.2 cells, an immortalised mouse neural progenitor cell line derived from neonatal mouse cerebellum. This is consistent with N-truncated Shh release from cholesterol-depleted HeLa cells, Panc1 cells and MIA PaCa-2 cells (Manikowski et al., 2019) and demonstrates that proteolytic Shh processing during release is not limited to our experimental Bosc23 system.

Manikowski, D., P. Jakobs, H. Jboor, and K. Grobe. 2019. Soluble Heparin and Heparan Sulfate Glycosaminoglycans Interfere with Sonic Hedgehog Solubilization and Receptor Binding. *Molecules* **24**.

Wang, Q., D. E. Asarnow, K. Ding, R. K. Mann, J. Hatakeyama, Y. Zhang, Y. Ma, Y. Cheng, and P. A. Beachy. 2021. Dispatched uses Na(+) flux to power release of lipid-modified Hedgehog. *Nature* **599**: 320-24.

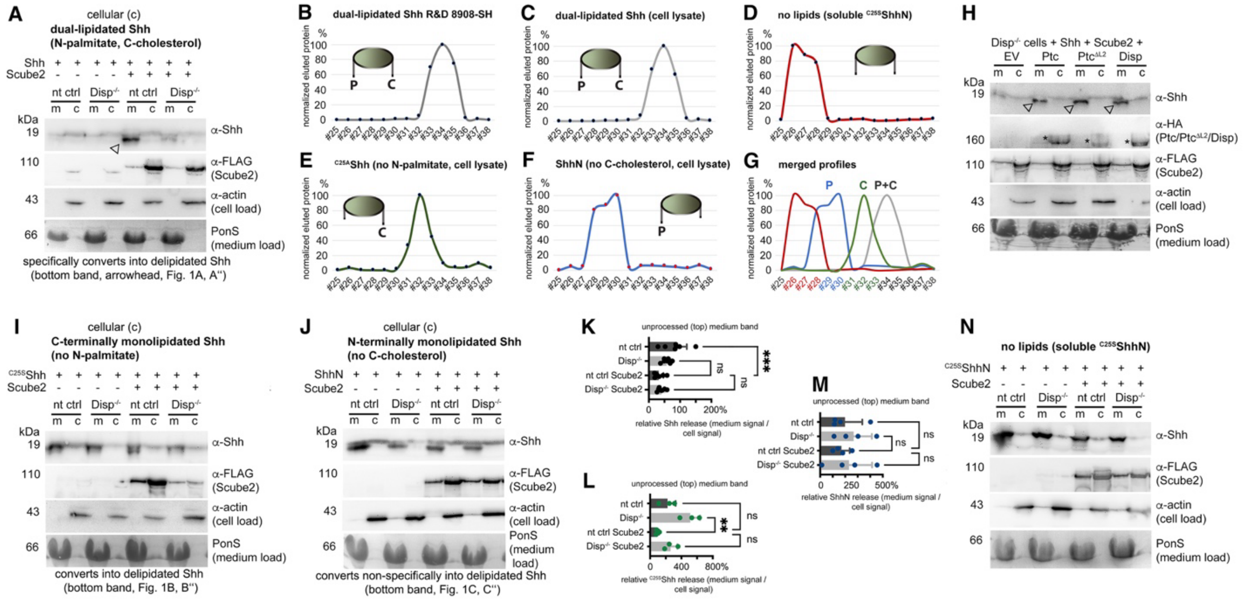


**Figure 1 - Figure Supplement 1. Loading controls, activity controls, and standards on the same stripped blots.** **A)** Actin served as a loading control for cell lysates and PonceauS as a loading control in media, and α-FLAG antibodies detected FLAG-tagged Scube2 on the same (stripped) blot. The arrowhead indicates Shh specifically solubilized by Scube2 and Disp. **B-G)** RP-HPLC of Shh variant standards. **B,C)** Elution profiles of commercial, dual-lipidated R&D 8908-SH Shh (B) and of dual-lipidated Shh from Bosc23 cell lysates (C) are similar. Cartoons of Shh proteins indicate the presence or absence of N-terminal palmitate (P) and C-terminal cholesterol (C). **D)** Bosc23-expressed non-lipidated control $^{C25A}$ShhN elutes in early fractions from the column. **E,F)** The elution profile of monolipidated $^{C25A}$Shh (E) partially overlaps with that of dual-lipidated Shh, and the elution profile of palmitoylated ShhN (F) partially overlaps with that of $^{C25A}$ShhN. The increased hydrophobicity of cholesteroylated $^{C25A}$Shh over palmitoylated ShhN is consistent with the finding that cholesterol membrane anchoring strength is similar to dual lipidation motifs such as double geranylgeranylation or palmitoylation plus farnesylation in other lipidated proteins (Peters et al., 2004). **G)** Merged profiles. Elution profiles are expressed relative to the highest protein amount in a given fraction (set to 100%). fr#: Fraction numbers are colored according to Shh lipidation status in this figure and in other figures. **H)** Co-expressed transgenic Ptch1, transgenic Ptch1$^{\Delta L2}$ lacking most of the second extracellular loop, and Disp-enhanced processed Shh release from Disp$^{-/-}$ cells into the media. Arrowheads indicate Shh specifically solubilized by Scube2 and Disp or Ptch1. Empty vector (EV)-transfected Disp$^{-/-}$ cells served as negative controls. **I,J)** Actin served as a loading control for cell lysates and PonceauS as a loading control in media, and α-FLAG antibodies detected FLAG-tagged Scube2. **K-M)** Quantification of the lower electrophoretic mobility band (the top band) of Shh (A, K), $^{C25S}$Shh (I, L), and ShhN (J, M) released from nt ctrl and Disp$^{-/-}$ cells. Amounts of solubilized and cellular proteins on immunoblots were determined and expressed as % increase or decrease relative to the respective cellular fraction, which was always set to 100%. ***: p=0.0005, **: p=0.0014, n.s.: p>0.05. **N)** Actin served as a loading control for cell lysates and PonceauS as a loading control in media, and α-FLAG antibodies detected FLAG-tagged Scube2, as always on the same (stripped) blot.

Peters, C., A. Wolf, M. Wagner, J. Kuhlmann, and H. Waldmann. 2004. The cholesterol membrane anchor of the Hedgehog protein confers stable membrane association to lipid-modified proteins. *Proc Natl Acad Sci U S A* **101**: 8531-6.

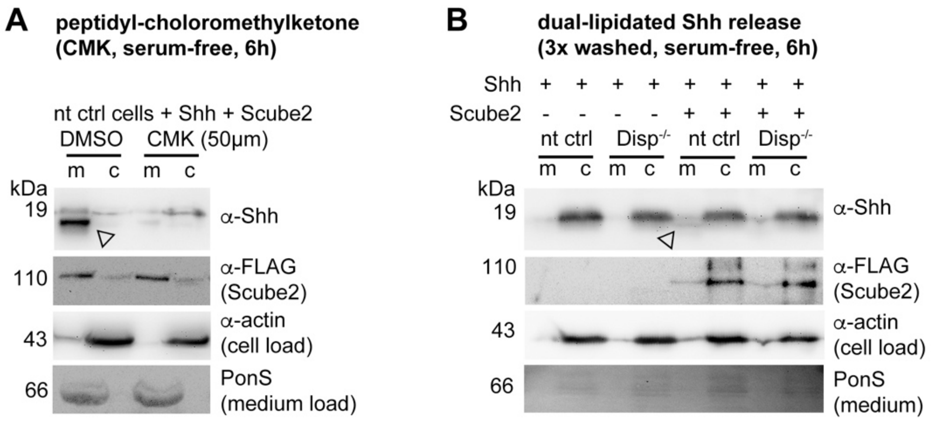


**Figure 2 - Figure Supplement 1. Loading controls. A,B)** Actin, PonceauS, and FLAG-tagged Scube2 loading controls for (the same stripped) blots shown in Fig. 2.

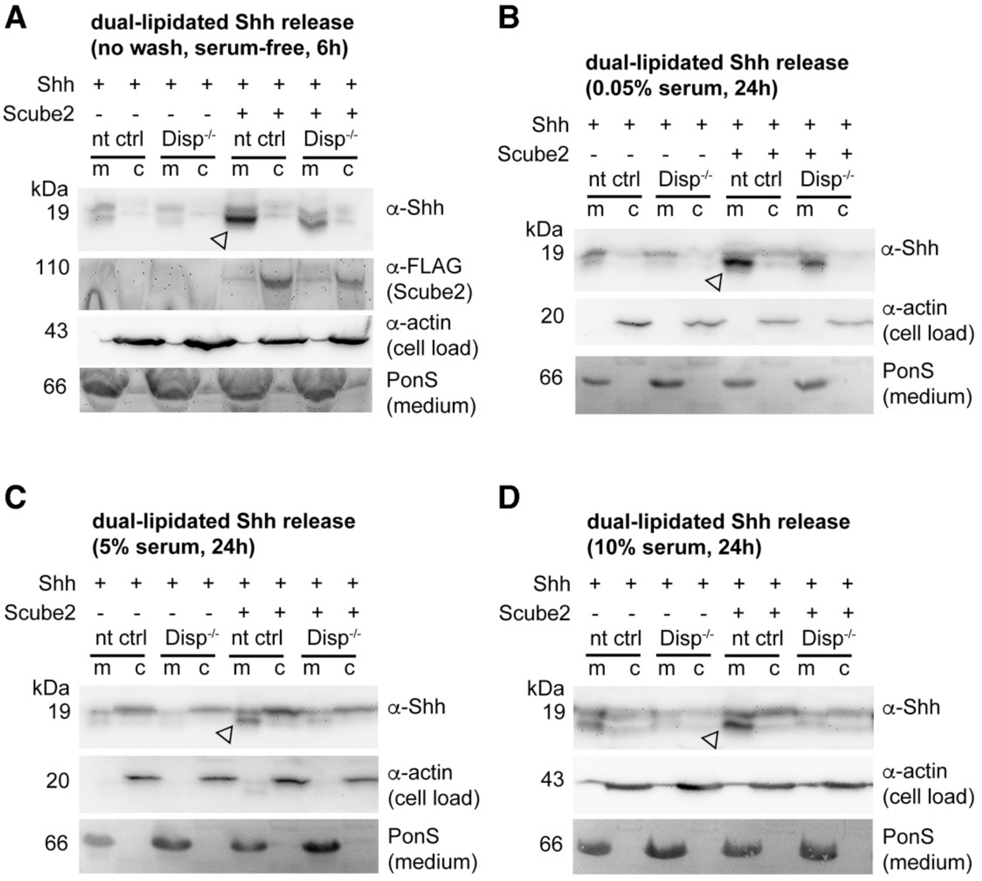


**Figure 3 - Figure Supplement 1. Loading controls. A-D)** Actin, PonceauS, and FLAG-tagged Scube2 loading controls for blots shown in Fig. 3. Arrowheads indicate Shh specifically solubilized by Disp and Scube2.

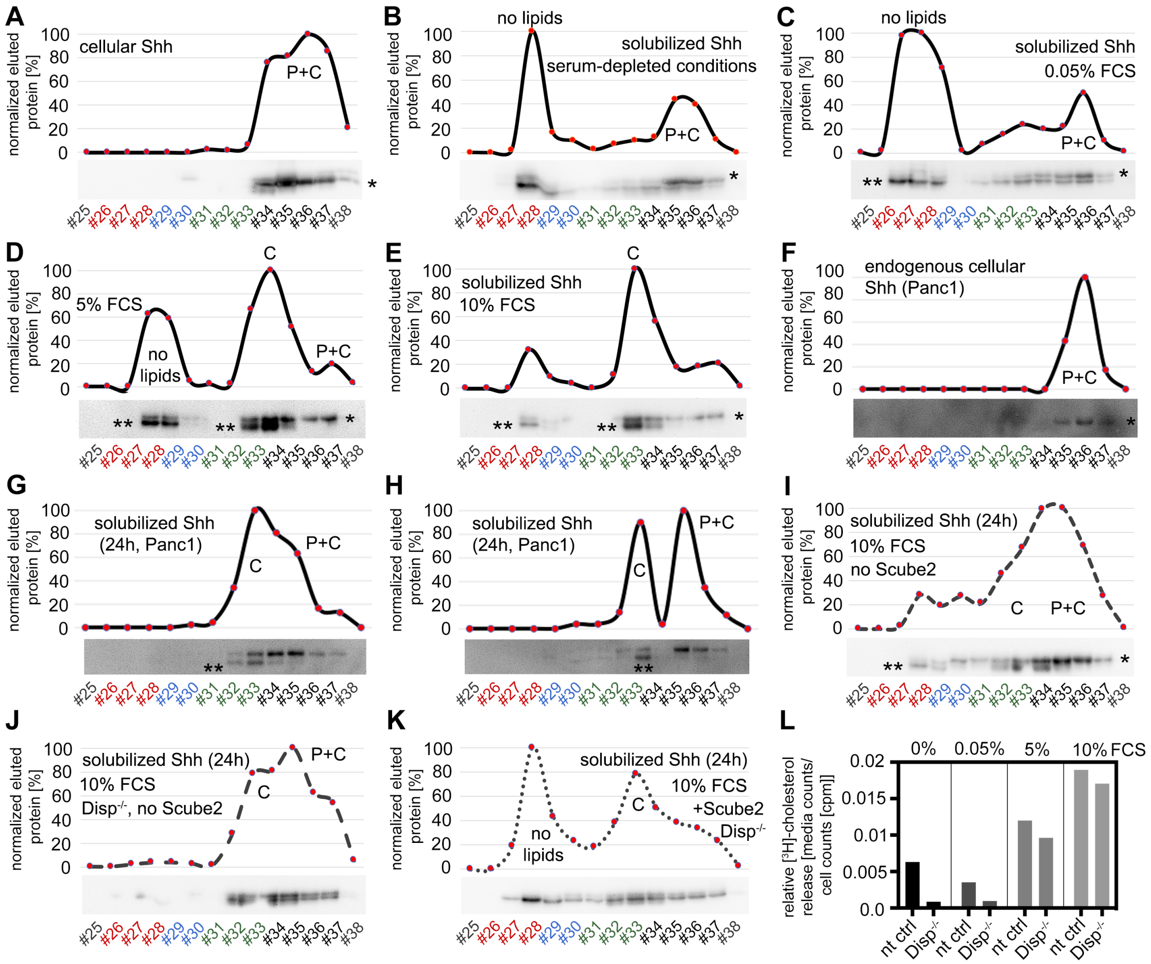


**Figure 3 – Figure Supplement 2. Source data and RP-HPLC profiles. A)** Dual-lipidated control Shh in cell lysates ("top" band, indicated by *). **B-E)** Serum modulates the conversion of dual-lipidated Shh into variably lipidated soluble forms. Shh was enriched by heparin-agarose pulldown before analysis. Bottom bands (** in the blot image) represent processed proteins derived from the dually delipidated Shh precursors (corresponding to the non-lipidated control $^{C25A}$ShhN, Fig. S2) or the cholesteroylated N-truncated form (fractions #32-34). Variable amounts of dual-lipidated proteins are also detected (top right bands (*), fractions #35-37). P: bound palmitate, C: bound cholesterol, P+C: dual-lipidated proteins. **F-H)** Serum modulates the conversion of endogenous dual-lipidated Shh (F) into mono-lipidated soluble forms (lower bands, marked with ** in the blot images) in two independent analyses (G,H). **I-K)** Increased relative amounts of dual-lipidated Shh released under physiologically irrelevant conditions. Proteins were released for 24 h in DMEM+10% FCS to compensate for impaired Shh solubilizations due to the absence of Scube2 and Disp. **I)** Lack of Scube2 in Disp-expressing cells decreases delipidated Shh solubilization (**) and monolipidated Shh release into the medium and increases the relative amounts of dual-lipidated Shh (*). **J)** Scube2 and Disp deficiency reduces delipidated and monolipidated Shh release, making dual-lipidated Shh the major soluble fraction detectable after 24h release. **K)** Scube2 restores some delipidated Shh release from Disp$^{-/-}$ cells. **L)** Bosc23 cells were loaded with [$^{3}$H]-cholesterol for 48 h and washed, and cholesterol release was measured for 3 h into serum-depleted media or medium with 0.05%, 5%, or 10% FCS. [$^{3}$H]-cholesterol signals in media were normalized with cellular counts. Note that cholesterol export from Disp$^{-/-}$ cells was always reduced if compared to nt ctrl cells under the same experimental conditions. n=2 for all measurements.

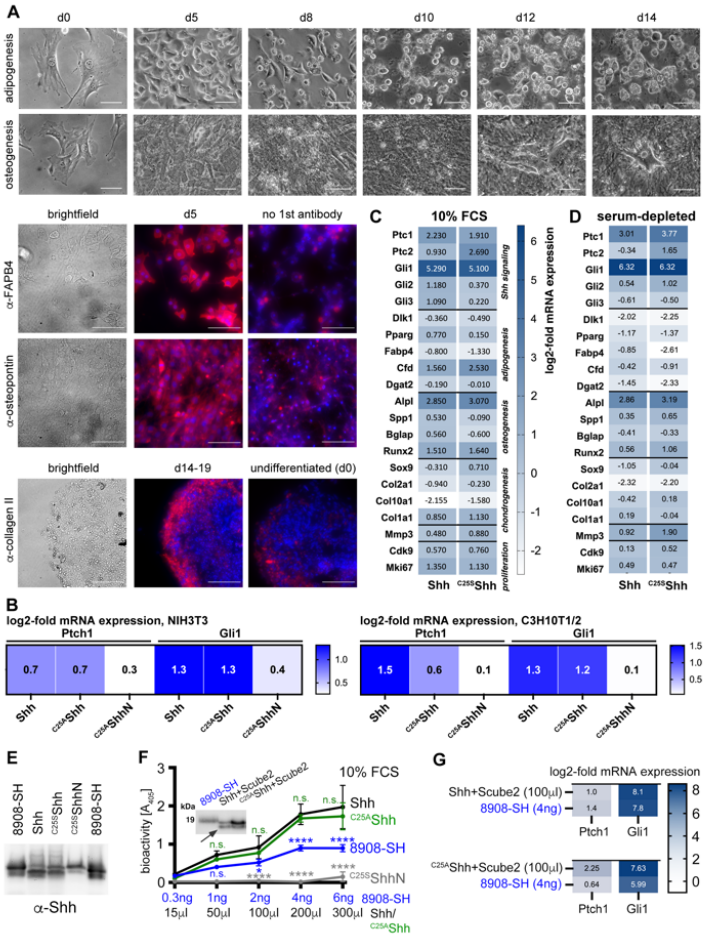


**Figure Supplement 5. Confirmed multipotency of C3H10T1/2 cells and similar activities of palmitoylated and non-palmitoylated Shh released into serum containing media. A)** C3H10T1/2 cells were cultured for the indicated time periods and their specific differentiation into adipocytes, osteoblasts or chondrocytes was induced. Differentiation was confirmed by morphology (top) in combination with lipid droplet staining with OilRed to confirm adipogenesis (not shown), alizarin red staining to identify $Ca^{2+}$-containing osteocytes (not shown), by anti-FABP-4 antibody binding that confirms adipogenic differentiation l, anti-collagen II antibody binding to confirm chondrogenic differentiation, and anti-osteopontin antibody binding to confirm osteogenesis. Cell nuclei were counterstained with DAPI (blue). Scale bars: 100 µm. **B)** N-processed soluble Shh and non-palmitoylated $^{C25A}$Shh induced similar increases in Ptch1 and Gli1 mRNA expression in stimulated NIH3T3 cells and C3H10T1/2 osteoblast progenitor cells. Non-lipidated $^{C25A}$ShhN served as an inactive control. Gene expression was normalized to mock-treated cells. **C,D)** N-processed Shh and partially N-processed $^{C25S}$Shh induced similar gene expression profiles when released into 10% FCS-containing DMEM or into serum-free DMEM supplemented with 300µg/ml MβCD (D). MβCD served as an alternative chaperone for the Shh C-cholesterol to show that associated serum factors (such as LPPs) do not alter the Shh activity profiles. See Tables S2 and S3 for qPCR target gene information. **E)** Normalization of soluble proteins used for the assays shown in Fig. 4B. **F,G)** Activities of dual-lipidated Shh and depalmitoylated Shh variants solubilized into serum-containing media. **F)** Shh, $^{C25A}$Shh and $^{C25S}$ShhN were solubilized from Scube2- and Disp expressing cells into media containing 10% FCS. Inset: Immunoblotted proteins were used for quantification and visualization of dually lipidated R&D 8908-SH, Shh, $^{C25A}$Shh and unlipidated $^{C25S}$ShhN. The arrow indicates proteolytic processing of the palmitoylated N-terminal peptide from Shh and $^{C25A}$Shh, but not from R&D 8908-SH. Graph: 0-4 ng of R&D 8908-SH dissolved into DMEM+10% FCS induced C3H10T1/2 reporter differentiation in a concentration-dependent manner, as determined by Alp activity measured at 405nm (blue line). Similar amounts of solubilized Shh, as determined by immunoblotting, increased C3H10T1/2 osteogenesis over that of the dual-lipidated R&D 8908-SH standard (black line). $^{C25A}$Shh showed similarly high bioactivity after release into serum-containing medium (green line). Artificial unlipidated soluble $^{C25S}$ShhN showed low activity (gray line). Data were normalized to mock-treated C3H10T1/2 cells. One-way ANOVA, Sidaks multiple comparisons test. ****: p>0.0001, *: p= 0.03, n.s.: p>0.05. **G)** Similar induction of transcription of the Hh target genes Ptch1 and Gli1 in C3H10T1/2 reporter cells. Cells were stimulated with 4ng R&D 8908-SH or similar amounts of Shh and $^{C25A}$Shh. Shh and $^{C25A}$Shh were both solubilized from Bosc23 cells into serum containing media in the presence of Scube2.

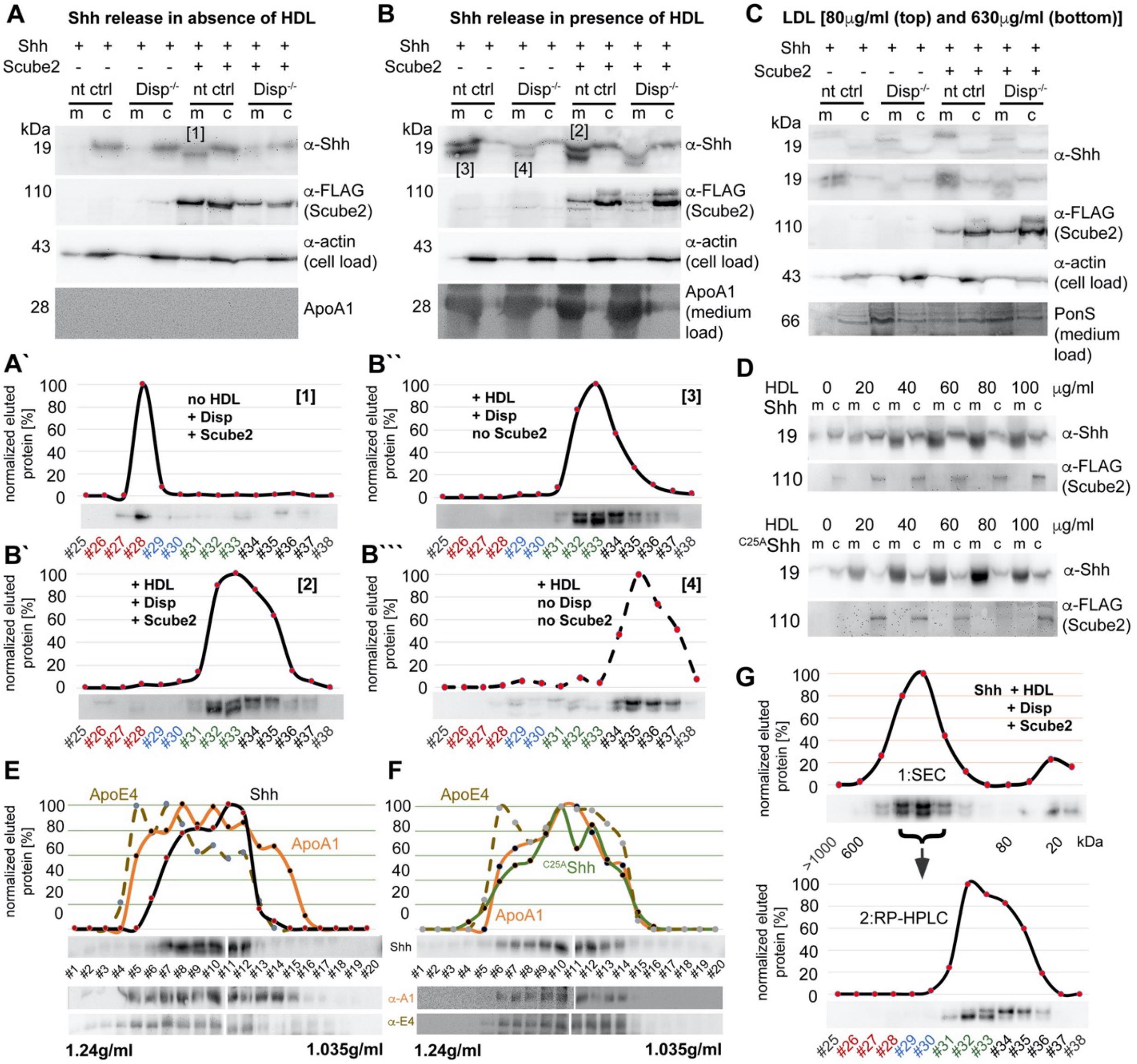


**Figure 5 – Figure Supplement 1. Loading controls. A,B)** Actin and FLAG-tagged Scube2 loading controls for the blots shown in Fig. 2. No residual BSA was detected in these media as a result of repeated careful washing of the cells prior to the addition of DMEM+HDL. ApoA1 was therefore used as the loading control. **A'-B''')** RP-HPLC profiles and source data of aliquots of samples [1-4] as shown in A, B. Note that Disp$^{-/-}$ cells release high relative amounts of dual-lipidated Shh in the absence of Scube2 (B'''). This suggests that the transfer of dual-lipidated Shh to HDL is unlikely to be physiologically relevant. In contrast, Disp and Scube2 specifically enhance the release of Shh in delipidated (A') or monolipidated (B',B'') form, depending on the presence or absence of HDL. **C)** LDL does not increase Shh solubilization from Disp-expressing cells. Transfection and loading controls are shown for the stripped top blot (80μg/ml LDL). **D)** In contrast, HDL increases the release of Shh and of cholesteroylated, non-palmitoylated $^{C25A}$Shh in a concentration-dependent manner. **E, F)** Shh (E) and $^{C25A}$Shh (F) were transfected into Disp-expressing Bosc23 cells and released overnight in the presence of 40mg/ml HDL in 1.2ml DMEM. Density gradient centrifugation of the solubilized material supports the physical interaction of both proteins with ApoA1 and ApoE4-bearing HDL of known corresponding density (1.1-1.2 g/ml). **G)** Shh released from transfected Disp-expressing cells was released in the presence of HDL and subjected to SEC. High molecular weight SEC fractions were subsequently analyzed by RP-HPLC, demonstrating that all HDL-associated proteins are cholesterylated.

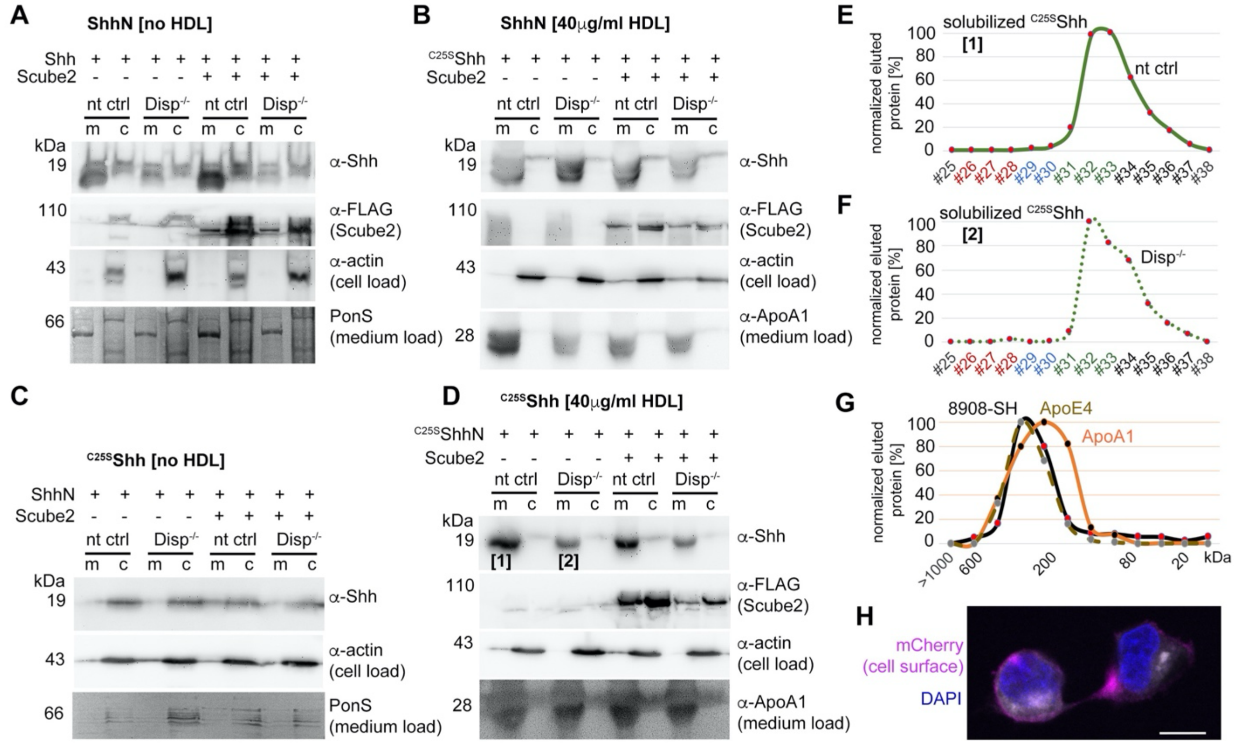


**Figure 7 – Figure Supplement 1. Loading controls. A-D)** Actin, PonceauS, and FLAG-tagged Scube2 loading controls for the blots shown in Fig. 6. A,B. **E,F)** RP-HPLC profiles of samples [1] and [2] shown in D. Note that proteins released from nt ctrl cells and Disp$^{-/-}$ cells are both cholesterylated. This suggests that overexpressed monolipidated Shh is only loosely associated with the plasma membrane and can "leak out", or desorb, and thereby associate with HDL in a non-enzymatic manner. **G)** Purified commercial 8908-SH Shh and HDL were mixed, incubated for 10 min, and analyzed by SEC. This revealed rapid spontaneous association and explains the background desorption of dual-lipidated Shh in our cellular assays. Note the excellent 8908-SH Shh co-elution with the ApoE4-bearing HDL fraction, suggesting that this mobile apolipoprotein may have facilitated spontaneous 8908-SH Shh association, or that other properties of ApoE-bearing "late" HDL (Sacks et al., 2018) somehow facilitate Shh association. **H)** Dual-lipidated mCherry is efficiently secreted to the outer plasma membrane leaflet of producing Bosc23 cells. Intracellular mCherry fluorescence is shown in white, α-mCherry antibody binding to the surface of non-permeabilized cells is shown in margenta, and DAPI staining of the nucleus is shown in blue. Scale bar: 10 µm.

Sacks, F. M., and M. K. Jensen. 2018. From High-Density Lipoprotein Cholesterol to Measurements of Function: Prospects for the Development of Tests for High-Density Lipoprotein Functionality in Cardiovascular Disease. *Arterioscler Thromb Vasc Biol* **38**: 487-99.

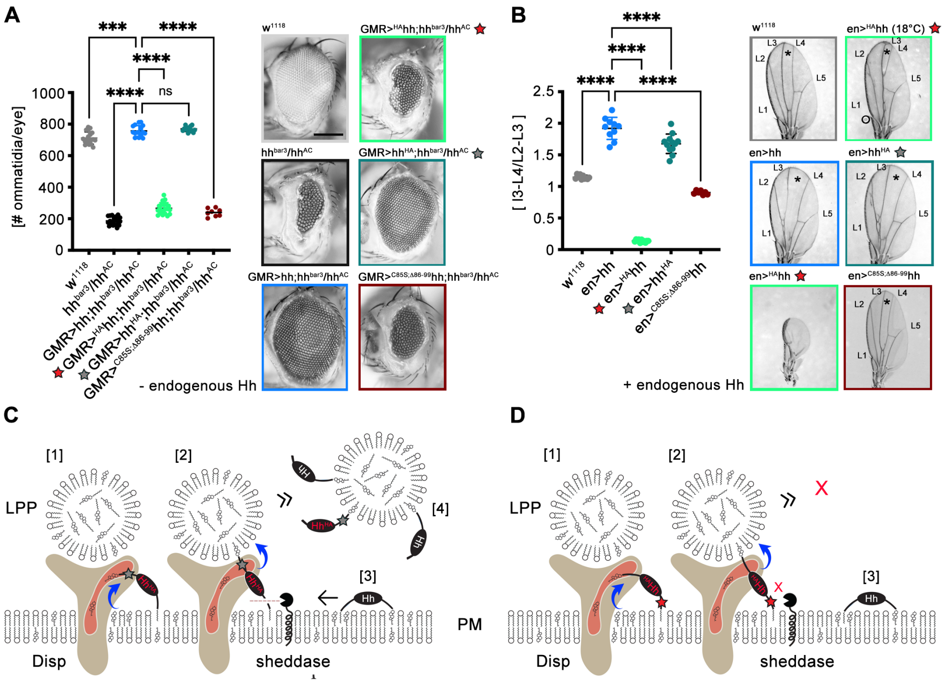


**Figure 8 – Figure Supplement 1. *In vivo* support for N-terminal shedding during Disp-mediated Hh export.** Experiments were independently repeated from established fly stocks and from published protocols to demonstrate that N-terminal, but not C-terminal proteolytic Hh processing is required during release (Schürmann et al., 2018; Kastl et al., 2018; Manikowski et al., 2021). **A)** Wild-type flies (w$^{1118}$) were used as positive controls. Eye discs lacking most Hh expression (in an $hh^{bar3}$/hh$^{AC}$ background) develop into small eyes, and eye-disc-specific *GMR-GAL4*-controlled hh or hh$^{HA}$ (Here, a protease-resistant hemagglutinin tag was inserted into the cholesteroylated C-terminal Hh peptide, gray star) restore eye development in the $hh^{bar3}/hh^{AC}$ background. In contrast, *GMR-GAL4*-controlled $^{HA}$hh expression (where the hemagglutinin tag was inserted into the N-palmitoylated Hh peptide to block sheddase function, red star) in the same background does not restore eye development. A non-palmitoylated N-truncated control variant (GMR>$^{C85S;\Delta N86\text{-}99}$hh) is also inactive. Left: Quantification of phenotypes, right: representative examples of adult compound eyes. Scale bar: 100 µm. One-way ANOVA, Dunnett's multiple comparisons test. ****: $p<0.0001$, ***: $p<0.001$, n.s.: not significant ($p>0.05$). **B)** The same Hh transgenes were produced in the posterior compartment of the wing disc pouch at 25°C under the control of *en-GAL4*. Cells in this compartment also produce endogenous Hh under the control of *en*, resulting in the simultaneous presence of endogenous and transgenic Hh at the cell surface. In adult wild-type flies (w$^{1118}$) expressing only endogenous Hh, the wing disc develops into a wing blade with five longitudinal veins (L1-5) (Hartl et al., 2014). The central L3-L4 intervein tissue (asterisk) is derived from cells that receive the highest dose of Hh and thus serves as a readout for Hh function in the developing disc. As a consequence, *en-GAL4*-controlled overexpression of hh or hh$^{HA}$ leads to an expansion of the L3-L4 intervein area. In striking contrast, wing formation is abolished by overexpression of transgenic $^{HA}$hh at 25°C under the control of *en-GAL4*, despite continued expression of endogenous hh. Reduced Gal4 promotor function at 18°C largely restores wing development with the exception of the most Hh-dependent L3-L4 domain. This indicates that overexpressed $^{HA}$hh, but not hh$^{HA}$, dominant-negatively inhibits endogenous Hh signaling in the developing wing, most likely by blocking N-terminal shedding. The inactive control mutant $^{C85S;\Delta N86\text{-}99}$hh does not suppress endogenous Hh function. One-way ANOVA, Dunnett's multiple comparisons test. ****: $p<0.0001$. See Table S1 for source data. **C)** We suggest that C-terminal protein tags (indicated by the gray star) do not interfere with Disp- and cholesterol-dependent protein extraction [1] and transfer [2] to LPPs. As a consequence, extraction and transfer of endogenous Hh [3] to the same LPPs [4] is not impaired. **D)** In contrast, impaired N-processing by the tag (red star) may stall Disp-mediated protein transfer [1,2] and create a bottleneck for endogenous Hh export [3], which in turn also reduces endogenous Hh signaling in the disc as well.

Hartl, T. A., and M. P. Scott. 2014. Wing tips: The wing disc as a platform for studying Hedgehog signaling. *Methods* **68**: 199-206.

Kastl, P., D. Manikowski, G. Steffes, S. Schurmann, S. Bandari, C. Klambt, and K. Grobe. 2018. Disrupting Hedgehog Cardin-Weintraub sequence and positioning changes cellular differentiation and compartmentalization in vivo. *Development* **145**: dev167221.

Manikowski, Dominique, Philipp Kastl, Sabine Schürmann, Kristina Ehring, Georg Steffes, Petra Jakobs, and Kay Grobe. 2021. C-Terminal Peptide Modifications Reveal Direct and Indirect Roles of Hedgehog Morphogen Cholesteroylation. *Frontiers in Cell and Developmental Biology* **8**.

Schürmann, S., G. Steffes, D. Manikowski, P. Kastl, U. Malkus, S. Bandari, S. Ohlig, C. Ortmann, R. Rebollido-Rios, M. Otto, H. Nusse, D. Hoffmann, C. Klämbt, M. Galic, J. Klingauf, and K. Grobe. 2018. Proteolytic processing of palmitoylated Hedgehog peptides specifies the 3-4 intervein region of the Drosophila wing. *Elife* **7**: e33033.

**Table Supplement 1. Raw data used for the figures in this study.**

| Figure | Cell type | Constructs used/treatment | Values (Mean)±SD, n | P-values |
|---|---|---|---|---|
| 1A` | nt | Shh±Scube2 | 48.7±11% / 325±46%, n=7 | <0.0001 |
| 1A` | Disp$^{-/-}$ / nt | Shh±Scube2 | 31.9±17% / 325±46%, n=7 | <0.0001 |
| 1A` | Disp$^{-/-}$ / nt | Shh+Scube2 | 74.9±26% / 325±46%, n=7 | <0.0001 |
| 1B` | nt | $^{C25S}$Shh±Scube2 | 207±311% / 314±248%, n=3 | 0.8503 |
| 1B` | Disp$^{-/-}$ / nt | $^{C25S}$Shh±Scube2 | 39.7±28% / 314±248%, n=3 | 0.2863 |
| 1B` | Disp$^{-/-}$ / nt | $^{C25S}$Shh+Scube2 | 31.7±29% / 314±248%, n=3 | 0.2679 |
| 1C` | nt | ShhN±Scube2 | 810±131% / 607±310%, n=4 | 0.5065 |
| 1C` | Disp$^{-/-}$ / nt | ShhN±Scube2 | 960±300% / 607±310%, n=4 | 0.1366 |
| 1C` | Disp$^{-/-}$ / nt | ShhN+Scube2 | 194±148% / 607±310%, n=4 | 0.0739 |
| 1D` | nt | $^{C25S}$ShhN±Scube2 | 1048±21% / 1003±411%, n=3 | 0.9913 |
| 1D` | Disp$^{-/-}$ / nt | $^{C25S}$ShhN±Scube2 | 894±193% / 1003±411%, n=3 | 0.9033 |
| 1D` | Disp$^{-/-}$ / nt | $^{C25S}$ShhN+Scube2 | 801±177% / 1003±411%, n=3 | 0.6345 |
| 2C | Bosc23 | CMK/DMSO | 77%±38% / 501%±249%, n=6 | 0.0021 |
| 2E | nt | Shh±Scube2 | 33.3±18% / 62±21%, n=6 | 0.0216 |
| 2E | Disp$^{-/-}$ / nt | Shh±Scube2 | 27.7±14.5% / 62±21%, n=6 | 0.0059 |
| 2E | Disp$^{-/-}$ / nt | Shh+Scube2 | 51±13% / 62±21%, n=6 | 0.5483 |
| 3B` | nt | Shh±Scube2 .05% FCS | 87.3±64% / 377±91%, n=3 | 0.0013 |
| 3B` | Disp$^{-/-}$ / nt | Shh±Scube2 .05% FCS | 62±15% / 377±91%, n=3 | 0.0008 |
| 3B` | Disp$^{-/-}$ / nt | Shh+Scube2 .05% FCS | 92±59% / 377±91%, n=3 | 0.0015 |
| 3C` | nt | Shh±Scube2 5% FCS | 92±50% / 299±112%, n=3 | 0.0158 |
| 3C` | Disp$^{-/-}$ / nt | Shh±Scube2 5% FCS | 78±45% / 299±112%, n=3 | 0.0112 |
| 3C` | Disp$^{-/-}$ / nt | Shh+Scube2 5% FCS | 67±44% / 299±112%, n=3 | 0.0087 |
| 3D` | nt | Shh±Scube2 10% FCS | 52±45% / 338±77%, n=3 | 0.0001 |
| 3D` | Disp$^{-/-}$ / nt | Shh±Scube2 10% FCS | 30±12% / 338±77%, n=3 | <0.0001 |
| 3D` | Disp$^{-/-}$ / nt | Shh+Scube2 10% FCS | 55±12% / 338±77%, n=3 | 0.0002 |
| 3E | nt | Shh±Scube2 10% FCS | 152±12% / 158±59%, n=3 | 0.9916 |
| 3E | Disp$^{-/-}$ / nt | Shh±Scube2 10% FCS | 112±3% / 158±59%, n=3 | 0.2659 |
| 3E | Disp$^{-/-}$ / nt | Shh+Scube2 10% FCS | 105±26% / 158±59%, n=3 | 0.1841 |
| 4A | C3H10T1/2 | mock/Shh (125µl) | 0.13±0.02au / 0.877±0.14 au, n=6 | <0.0001 |
| 4A | C3H10T1/2 | Shh/ $^{C25S}$Shh (125µl) | 0.877±0.14 au / 0.872±0.09au, n=6 | >0.9999 |
| 4A | C3H10T1/2 | Shh/$^{C25S}$Shh (250µl) | 2.16±0.1au / 2.69±0.28au, n=6 | <0.0001 |
| 4B | C3H10T1/2 | Shh/ $^{C25S}$Shh (1x) Ptch1 | 2.66±0.15 / 2.64±0.08, n=3 | >0.9999 |
| 4B | C3H10T1/2 | Shh/ $^{C25S}$Shh (2x) Ptch1 | 3.11±0.49 / 3.31±1, n=3 | 0.9938 |
| 4B | C3H10T1/2 | Shh/ $^{C25S}$Shh (1x) Gli1 | 5.7±0.6 / 5.33±0.26, n=3 | 0.9446 |
| 4B | C3H10T1/2 | Shh/ $^{C25S}$Shh (2x) Gli1 | 5.5±0.79 / 5.5±1.14, n=3 | >0.9999 |
| 4B | C3H10T1/2 | Shh (1x) / $^{C25S}$ShhN (2x) Ptch1 | 2.66±0.15 / 0.78±0.48, n=3 | 0.0083 |
| 4B | C3H10T1/2 | $^{C25S}$Shh (1x) / $^{C25S}$ShhN (2x) Ptch1 | 2.64±0.08 / 0.78±0.48, n=3 | 0.009 |

| | | | | |
|---|---|---|---|---|
| 4B | C3H10T1/2 | Shh (2x) / $^{C25S}$ShhN (2x) Ptch1 | 3.11±0.49 / 0.78±0.48, n=3 | 0.0019 |
| 4B | C3H10T1/2 | $^{C25S}$Shh (2x) / $^{C25S}$ShhN (2x) Ptch1 | 3.31±1 / 0.78±0.48, n=3 | 0.0011 |
| 4B | C3H10T1/2 | Shh (1x) / $^{C25S}$ShhN (2x) Gli1 | 5.7±0.6 / 1.84±0.68, n=3 | 0.0003 |
| 4B | C3H10T1/2 | $^{C25S}$Shh (1x) / $^{C25S}$ShhN (2x) Gli1 | 5.33±0.26 / 1.84±0.68, n=3 | 0.0007 |
| 4B | C3H10T1/2 | Shh (2x) / $^{C25S}$ShhN (2x) Gli1 | 5.5±0.79 / 1.84±0.68, n=3 | 0.0005 |
| 4B | C3H10T1/2 | $^{C25S}$Shh (2x) / $^{C25S}$ShhN (2x) Gli1 | 5.5±1.14 / 1.84±0.68, n=3 | 0.0005 |

| | | | | |
|---|---|---|---|---|
| 5C | nt | Shh±Scube2 | 446%±177% / 392%±176%, n=7 | 0.7727 |
| 5C | Disp$^{-/-}$ / nt | Shh±Scube2 | 131%±35% / 392%±176%, n=7 | 0.0023 |
| 5C | Disp$^{-/-}$ / nt | Shh+Scube2 | 101%±38% / 392%±176%, n=7 | 0.0008 |

| | | | | |
|---|---|---|---|---|
| 6B | C3H10T1/2 | Shh/ $^{C25S}$Shh (1x) Ptch1 | 1.685±0.31 / 1.44±0.16, n=3 | 0.9632 |
| 6B | C3H10T1/2 | Shh/ $^{C25S}$Shh (2x) Ptch1 | 1.653±0.49 / 1.89±0.33, n=3 | 0.9683 |
| 6B | C3H10T1/2 | Shh/ $^{C25S}$Shh (1x) Gli1 | 4.24±0.23 / 4.47±0.26, n=3 | 0.9691 |
| 6B | C3H10T1/2 | Shh/ $^{C25S}$Shh (2x) Gli1 | 4.44±0.74 / 4.47±1.03, n=3 | >0.9999 |
| 6B | C3H10T1/2 | Shh (1x) / $^{C25S}$ShhN (2x) Ptch1 | 1.685±0.31 / 1.3±0.6, n=3 | 0.0013 |
| 6B | C3H10T1/2 | $^{C25S}$Shh (1x) / $^{C25S}$ShhN (2x) Ptch1 | 1.44±0.16 / 1.3±0.6, n=3 | 0.0053 |
| 6B | C3H10T1/2 | Shh (2x) / $^{C25S}$ShhN (2x) Ptch1 | 1.653±0.49 / 1.3±0.6, n=3 | 0.0015 |
| 6B | C3H10T1/2 | $^{C25S}$Shh (2x) / $^{C25S}$ShhN (2x) Ptch1 | 1.89±0.33 / 1.3±0.6, n=3 | 0.0004 |
| 6B | C3H10T1/2 | Shh (1x) / $^{C25S}$ShhN (2x) Gli1 | 4.24±0.23 / 1.3±0.6, n=3 | 0.0008 |
| 6B | C3H10T1/2 | $^{C25S}$Shh (1x) / $^{C25S}$ShhN (2x) Gli1 | 4.47±0.26 / 1.3±0.6, n=3 | 0.0005 |
| 6B | C3H10T1/2 | Shh (2x) / $^{C25S}$ShhN (2x) Gli1 | 4.44±0.74 / 1.3±0.6, n=3 | 0.0005 |
| 6B | C3H10T1/2 | $^{C25S}$Shh (2x) / $^{C25S}$ShhN (2x) Gli1 | 4.47±1.03 / 1.3±0.6, n=3 | 0.0005 |

Figure Supplements

| | | | | |
|---|---|---|---|---|
| FS1B | Disp$^{-/-}$ / nt | Shh-NanoLuc | 65%±30% / 2745%±1186%, n=3 | <0.0173 |

| | | | | |
|---|---|---|---|---|
| F1S1K | nt | Shh±Scube2 | 83.7±38% / 31.5±15%, n=7 | 0.0005 |
| F1S1K | Disp$^{-/-}$ / nt | Shh±Scube2 | 56.9±13% / 31.5±15%, n=7 | 0.1034 |
| F1S1K | Disp$^{-/-}$ / nt | Shh+Scube2 | 40±11% / 31.5±15%, n=7 | 0.8103 |
| F1S1L | nt | $^{C25S}$Shh±Scube2 | 223±102% / 93.3±22%, n=3 | 0.2739 |
| F1S1L | Disp$^{-/-}$ / nt | $^{C25S}$Shh±Scube2 | 514±126% / 93.3±22%, n=3 | 0.0014 |

| | | | | |
|---|---|---|---|---|
| F1S1L | Disp$^{-/-}$ / nt | $^{C25S}$Shh+Scube2 | 262±87% / 93.3±22%, n=3 | 0.1307 |
| F1S1M | nt | ShhN±Scube2 | 199±137% / 170±66%, n=4 | 0.9819 |
| F1S1M | Disp$^{-/-}$ / nt | ShhN±Scube2 | 268±146% / 170±66%, n=4 | 0.6471 |
| F1S1M | Disp$^{-/-}$ / nt | ShhN+Scube2 | 230±185% / 170±66%, n=4 | 0.8752 |

| | | | | |
|---|---|---|---|---|
| FS5F | Bosc23 | Shh (50µl-300µl) | 0.7±0.1, 0.9±0.3, 1.8±0.27, 2.0±0.56, n=4 ea | |
| FS5F | Bosc23 | $^{C25S}$Shh (50µl-300µl) | 0.6±0.04, 0.77±0.02, 1.7±0.1, 1.7±0.35, n=4 ea | 0.89, 0.78, 0.91, 0.33 |
| FS5F | Bosc23 | $^{C25S}$ShhN (50µl-300µl) | 0.02±0.01, 0.04±0.01, 0.02±0.015, 0.15±0.12, n=4 ea | <0.0001 for all amounts |
| FS5F | HEK293 | R&D 8908-SH 1ng-6ng) | 0.4±0.04, 0.52±0.1, 0.9±0.08, 0.9±0.09, n=4 ea | 0.13, 0.033, <0.0001, <0.0001 |

| | | | | |
|---|---|---|---|---|
| F8S1A | eye disc | w$^{1118}$ control | 712±39, n=20 | 0.0002 |
| F8S1A | eye disc | | 186±26, n=20 | <0.0001 |
| F8S1A | eye disc | Hh | 757±34, n=16 | |
| F8S1A | eye disc | $^{HA}$Hh | 266±36, n=20 | <0.0001 |
| F8S1A | eye disc | Hh$^{HA}$ | 769±16, n=16 | 0.6999 |
| F8S1A | eye disc | $^{C85S;\Delta 86-99}$Hh | 239±25, n=8 | <0.0001 |
| F8S1B | wing disc | w$^{1118}$ control | 1.147±0.03, n=10 | <0.0001 |
| F8S1B | wing disc | Hh | 1.92±0.17, n=10 | |
| F8S1B | wing disc | $^{HA}$Hh | 0.14±0.02, n=11 | <0.0001 |
| F8S1B | wing disc | Hh$^{HA}$ | 1.68±0.15, n=12 | <0.0001 |
| F8S1B | wing disc | $^{C85S;\Delta 86-99}$Hh | 0.9±0.03, n=7 | <0.0001 |

**Table Supplement 2: qPCR target genes analyzed in this work**

| | | | | |
|---|---|---|---|---|
| 1 | control, normalization of 2-21 | **Actb** | β-actin | Control, cytoplasmic |
| 2 | Shh signaling | **Ptch1** | Patched homolog 1 | Receptor for Shh/Ihh/Dhh |
| 3 | Shh signaling | **Ptch2** | Patched homolog 2 | Possible receptor for Shh, plays a role in control of cellular growth |
| 4 | Shh signaling | **Gli1** | Zinc finger protein GLI1 | Transcriptional activator, binds to DNA consensus 5'-GACCACCCA-3', craniofacial/digit/CNS/gastrointestinal development. Direct target of Hh |
| 5 | Shh signaling | **Gli2** | Zinc finger protein GLI2 | Transcriptional activator (repressor) in Shh pathway, binds to DNA sequence 5'-GAACCACCCA-3' (part of TRE-2S regulatory element) |
| 6 | Shh signaling | **Gli3** | Zinc finger protein GLI3 | Dual function: transcriptional activator (GLI3A full length after phosphorylation and nuclear translocation) and repressor (GLI3R, C-terminally truncated) of Shh pathway, binds to 5'-GGGTGGTC-3' |
| 7 | osteogenic differentiation | **Alpl** | Alkaline phosphatase (pan-expressed) | Key role in skeletal mineralization by regulating levels of diphosphate (PPi) |
| 8 | osteogenic differentiation | **Spp1** | Secreted phosphoprotein 1 (= Osteopontin) | Binds to hydroxyapatite, part of mineralized matrix, important for cell-matrix interactions |
| 9 | osteogenic differentiation | **Bglap** | Bone γ-carboxy-glutamate protein (Osteocalcin) | Hormone, secreted by osteoblasts, functions in bone remodeling and energy metabolism |
| 10 | osteogenic differentiation | **Runx2** | Runt related transcription factor 2 | Essential for osteoblastic differentiation and skeletal morphogenesis |
| 11 | adipogenic differentiation | **Dlk1** | Delta like non-canonical Notch ligand 1 (= Pref1) | Preadipocyte marker. TM protein, inhibits adipocyte differentiation |
| 12 | adipogenic differentiation | **Ppar-γ** | Peroxisome proliferator-activated receptor γ | Proadipogenic. Ligand-activated TF, binds to DNA-specific PPAR response elements, key regulator of adipocyte differentiation and glucose homeostasis |
| 13 | adipogenic differentiation | **Fabp4** | Fatty acid-binding protein (=Ap2) | Differentiated adipocytes (lipid transport protein in adipocytes, binds long chain fatty acids and retinoic acid) |
| 14 | adipogenic differentiation | **Cfd** | Complement factor D (= adipsin) | Differentiated adipocytes |
| 15 | TAG biosynthesis | **Dgat2** | Diacylglycerin-acyltransferase 2 | Acylation of DAG, → triacylgylcerol (TAG) |

| | | | | |
|---|---|---|---|---|
| 16 | chondrogenic differentiation | **Sox9** | SRY (sex determining region Y)-bos 9 | Promotes expression of genes in chondrogenesis, including cartilage matrix protein-coding genes COL2A1, COL4A2, COL9A1, COL11A2 and ACAN, SOX5 and SOX6. |
| 17 | chondrogenic differentiation | **Col2α1** | Collagen type II α I | Fibril-forming, major component of cartilage |
| 18 | chondrogenic differentiation | **Col10α1** | Collagen type X α I | Product of hypertrophic condrocytes, localised to presumptive mineralisation zones of hyaline cartilage |
| 19 | chondrogenic differentiation | **Col1α1** | Collagen type I α I | Fibril-forming, most abundant protein of bone/skin/tendon ECM |
| 20 | chondrogenic differentiation | **Mmp3** | Matrix metallopeptidase 3 | Codes for Stromelysin-1, degrades fibronectin, laminin, gelatins of type I, III, IV, and V; collagens III, IV, X, and IX, and cartilage proteoglycans. Activates procollagenase. Promotes cartilage degeneration. |
| 21 | proliferation | **Cdk9** | Cyclin-dependent kinase 9 (CDC2-related kinase) | Involved in regulation of transcription, component of TAK/P-TEFb complex forms complex with/is regulated by CyclinT or CyclinK |
| 21 | proliferation | **Mki67** | Antigen identified by monoclonal antibody Ki67 | Required to maintain individual mitotic chromosomes (associates with surface), dispersed in the cytoplasm following nuclear envelope disassembly |

**Table Supplement 3: Information regarding amplicons and primers**

| name | isoform | accession number | FWD primer (5'-3') | REV primer (3'-5') | Amplicon length |
|---|---|---|---|---|---|
| **Actin β** | 1 | NM_007393.5 | CTATTGGCAACGAGCGGTTC | CGGATGTCAACGTCACACTTC | 124 |
| **Ptch1** | 2 | NM_001328514.1, NM_008957.3 | GGGCTACGACTATGTCTCTC | CTTTGATGAACCACCTCCAC | 99 |
| **Ptch2** | 2 | NM_001312903.1, NM_008958.3 | TCTGTGCCCTGCTTCTACTC | GCCCAGGAATCCCATGATAC | 101 |
| **Gli1** | 1 | NM_010296.2 | CCCTGGTGGCTTTCATCAAC | TGACTCATCTGAGGTGGGAATC | 109 |
| **Gli2** | 1 | NM_001081125.1 | CAACTCAGCAGCAGTAGCAG | CTCCGCTTATGAATGGTGATGG | 110 |
| **Gli3** | 1 | NM_008130.3 | GTGGTTCCTATGGGCACTTATC | GTCGGCTTAGGATCTGTTGATG | 108 |
| **Dlk1** | 4 | NM_001190703.1, NM_001190704.1, NM_001190705.1, NM_010052.5 | GGCTGTGTCAATGGAGTCTG | AAGCCCGAACGTCTATTTCG | 91 |
| **Pparg** | 4 | NM_001127330.2, NM_001308352.1, NM_001308354.1, NM_011146.3 | TCCATTCACAAGAGCTGACC | GGTGGAGATGCAGGTTCTAC | 99 |
| **Fabp4** | 1 | NM_024406.3 | GTGTGATGCCTTTGTGGGAAC | CATGCCTGCCACTTTCCTTG | 106 |
| **Cfd** | 3 | NM_001291915.2, NM_001329541.1, NM_013459.4 | CCTGAACCCTACAAGCGATG | CAACGAGGCATTCTGGGATAG | 117 |
| **Dgat2** | 1 | NM_026384.3 | GGCTGATAGCTGTGCTCTAC | GATGGGAAAGTAGTCTCGGAAG | 128 |
| **AlpI** | 5 | NM_001287172.1, NM_007431.3 | CTGCAAGGACATCGCATATCAG | CCACATCAGTTCTGTTCTTCGG | 104 |
| **Spp1** | 5 | NM_001204201.1, NM_001204202.1, NM_001204203.1, NM_001204233.1, NM_009263.3 | ACAGAATGCTGTGTCCTCTG | GGTCTCCATCGTCATCATCATC | 122 |
| **Bglap** | 2 | NM_001032298.3 (Bglap2), NM_007541.3 (Bglap) | CCAAGCAGGAGGGCAATAAG | CTCGTCACAAGCAGGGTTAAG | 122 |
| **Runx2** | 6 | NM_001145920.2, NM_001146038.2, NM_001271627.1, NM_001271630.1, NM_001271631.1, NM_009820.5 | ACACTGCCACCTCTGACTTC | GGGATGAAATGCTTGGGAACTG | 117 |
| **Sox9** | 1 | NM_011448.4 | CGGAACAGACTCACATCTCTCC | GACCCTGAGATTGCCCAGAG | 123 |
| **Col2a1** | 2 | NM_001113515.2, NM_031163.3 | CTGAAGGTGCTCAAGGTTCTC | GATCCTTTGGCTCCAGGAATAC | 109 |
| **Col10a1** | 1 | NM_009925.4 | TCTCCCAGCACCAGAATCTATC | CCATGAACCAGGGTCAAGAAC | 82 |
| **Col1a1** | 1 | NM_007742.4 | TGGTCCACAAGGTTTCCAAG | CATCTCCATTCTTGCCAGGAG | 107 |
| **Mmp3** | 1 | NM_010809.2 | ACTTGTCCCGTTTCCATCTC | GGTTCCAGAGAGTTAGACTTGG | 118 |

| | | | | | |
|---|---|---|---|---|---|
| **Cdk9** | 1 | NM_130860.3 | CAGCTCTGTGGCTCCATCAC | GTCCTTCACCTTCCGCTTCTG | 105 |
| **Mki67** | 1 | NM_001081117.2 | TGAGGCTGAGACATGGAGAC | GGTTCCTTTCCAAGGGACTTTC | 121 |
